# Establishing reliable blood biomarkers for trimethylamine N-oxide status in rodents: effects of oral choline challenge, dietary choline and fasting conditions

**DOI:** 10.1101/2024.12.06.627229

**Authors:** Ahmad Ud Din, Michael G. Sweet, Ashley M. McAmis, Juanita G. Ratliff, Anandh Babu Pon Velayutham, Andrew P. Neilson

## Abstract

Circulating concentrations of the gut microbial-mammalian metabolite trimethylamine N-oxide (TMAO) is linked to atherosclerosis risk. TMAO biosynthesis begins when dietary choline is converted to trimethylamine (TMA) by gut microbial TMA lyase. TMA is transported to the liver, where flavin-containing monooxygenases convert it to TMAO. While dietary modifications regulate TMAO production, the impact of different intake methods, including oral gavage, dietary supplementation, and conditions such as fasting versus non-fasting, has not been fully explored. Twelve female Sprague-Dawley rats were divided into three diet groups (*n* = 4 per group): no-choline (0% choline), low-choline (0.08% choline), and high-choline (1% choline). Choline and TMAO fasting and non-fasting blood concentrations, and their kinetics following an acute choline challenge, were assessed before and after a 2-week dietary intervention with the distinct choline dietary levels. Fasting choline was under tight control, with little effect of dietary choline. Non-fasting choline was more variable, with high dietary choline reflected in higher blood choline. Greater levels of dietary choline were reflected in significantly greater levels of TMAO, particularly for non-fasting levels. Kinetic profiling demonstrated additional information regarding the appearance and clearance of these compounds from blood. These results suggest that acute oral choline gavage is likely most suitable for studies targeting acute (direct) inhibitors, whereas a choline-rich diet with assessment of fasting and non-fasting blood levels is more suitable for studying alterations to TMAO production capacity. Future research should examine the impact on atherosclerosis biomarkers and microbiome diversity to deepen the understanding of TMAO regulation and its cardiovascular implications.

## 1. Introduction

Cardiovascular disease (CVD) is the leading cause of death worldwide and in the US [1,2]. According to a recent estimate, global deaths from CVD are 19.8 million annually [3]. Multiple risk factors are associated with CVD, such as poor diet, smoking, obesity, lack of physical activity, etc. The composition and function of the gut microbiome are also associated with various cardiovascular events [4] through the production of diet derived microbial metabolites [5]. Gut microbes can produce both beneficial and harmful metabolites, such as short-chain fatty acids and trimethylamine N-oxide (TMAO) [5,6], respectively.

Trimethylamine *N*-oxide (TMAO) was identified as a novel risk factor for CVD, specifically atherosclerosis, in 2011 by the Hazen group at the Cleveland Clinic [7,8]. Since then, elevated blood TMAO has been reported as a CVD and mortality risk factor [9,10], although controversy remains regarding its utility as a predictor of CVD risk [11,12]. TMAO is produced by sequential action of gut microbiota and the liver [8].

First, gut bacteria possessing choline trimethylamine (TMA) lyase enzymes (*cutC/D* gene cluster) metabolize choline into TMA [8]. TMA is then absorbed into the circulation and oxidized to TMAO by hepatic flavin-containing monooxygenase 3 (FMO3) [13]. Other quaternary amines, such as carnitine, can also be metabolized to TMA by similar microbial enzymes.

Choline is a semi-essential nutrient, and thus eliminating or drastically reducing choline intake is an unwise, impractical and ineffective strategy to reduce TMA and TMAO production [14,15]. Currently, there is no FDA-approved drug to manage elevated TMAO [16,17]. One strategy that has shown promise is direct inhibition of TMA lyases. For example, 3,3-dimethyl-1-butanol (DMB), a choline analog, inhibits TMA lyase and has shown a non-toxic potential to inhibit TMAO formation and prevent atherosclerosis *in vivo* [18,19]. However, due to the lack of approved pharmaceuticals to date, complementary non-drug strategies are needed. Dietary interventions are a logical strategy to reduce TMAO formation with the goal of preventing or slowing atherosclerosis/CVD [20,21]. Previous studies have demonstrated the cardioprotective roles of plant-based dietary interventions, including berries [20,22]. For example, blueberry supplementation mediates vascular inflammation, improves vascular dysfunction, and reduces circulating TMAO concentrations [23,24]. Polyphenols such as chlorogenic acid (CGA) are abundant in blueberries and have the potential to reduce choline microbial metabolism into TMA [25,26]. Chronic CGA supplementation improves the gut microbiome and protects against CVD in animal models [25,27]. It is currently unknown how CGA-rich foods may reduce TMA and TMAO, but it may act by acute mechanisms (such as direct TMA lyase inhibition) requiring the inhibitor and substrate to be present simultaneously, or mechanisms requiring chronic exposure, including shifts in microbiome taxonomic distribution, altered levels of *cutC/D*, and/or changes to FMO3 expression. The key step in lowering TMAO is reducing the conversion of choline to TMA by gut bacteria [18]. We recently developed an *ex vivo-in vitro* methodology to screen compounds for the ability to reduce microbial biotransformation of choline into TMA [28,29]. Using our fecal fermentation method, we tested the ability of CGA to inhibit TMA production [30]. We found CGA reduces both choline use and TMA production [30].

*In vivo* studies are needed to validate the efficacy of compounds identified *in vitro* as having potential TMA- and TMAO-lowering activities. There are various methods in place to use choline administration models either in a chronic diet or acute oral challenge (by gavage) [31,32]. However, there is a lack of consensus on the utility of various potential TMAO measures such as blood vs. urinary levels, or single time point vs. kinetics over time. There is no clearly accepted method to evaluate the efficacy of interventions that may reduce choline conversion to TMA and TMAO. Given that such interventions might work via acute or chronic intervention, experimental conditions such as blood collection in a fasting or non-fasting state, or use of chronic dietary choline vs. acute exposure, could yield vastly different results [33]. In the present study, we assessed the plasma TMAO responses with different experimental conditions such as acute oral choline gavage, dietary choline intervention, fasting, and non-fasting conditions. We found a clear difference in circulating TMAO between treatments of fasting, oral gavage, and dietary choline intervention in the preclinical model. This outcome could be considered for future in vivo studies to evaluate the efficacy of therapeutic compounds on circulating TMAO.

## 2. Materials and Methods

### 2.1 Reagents and chemicals

Choline chloride (C7017-5G), 3,3-dimethyl-1-butanol (183105-10G), ethyl 2-bromoacetate (133973-100G), ammonia (AM180-500MLGL), TMA (243205-100G), TMA-^13^C3-^15^N (730092), TMAO (317594-1G), and TMAO-d_9_ (DLM47791), were purchased from Sigma-Aldrich/Millipore (St. Louis, MO, USA). Ammonium formate (70221-25G-F), LC-MS grade acetonitrile (JT9829-3), and water were purchased from VWR International (Suwanee, GA, USA).

### 2.2 Animals

All animal experiments and procedures were approved by The North Carolina Research Campus (Kannapolis, NC) Institutional Animal Care and Use Committee (IACUC) under approval #20-017. The design of the study is shown in **Fig. 1**. A total of 12 female SAS Sprague-Dawley rats (age 7-8 weeks, 150-200g) were purchased from Envigo RMS (Indianapolis, IN, USA). On arrival, rats were kept in the Center for Laboratory Animal Science animal facility at the North Carolina Research Campus. Female rats are the best model of human FMO3 as they have higher FMO3 than males, allowing efficient conversion of TMA to TMAO [34–37]. Rats were caged in pairs and were allowed to assimilate for a week, provided with a standard chow (0.253% Choline; Teklad 8604, Envigo) diet, and access to tap water *ad libitum*.

**Figure 1.**
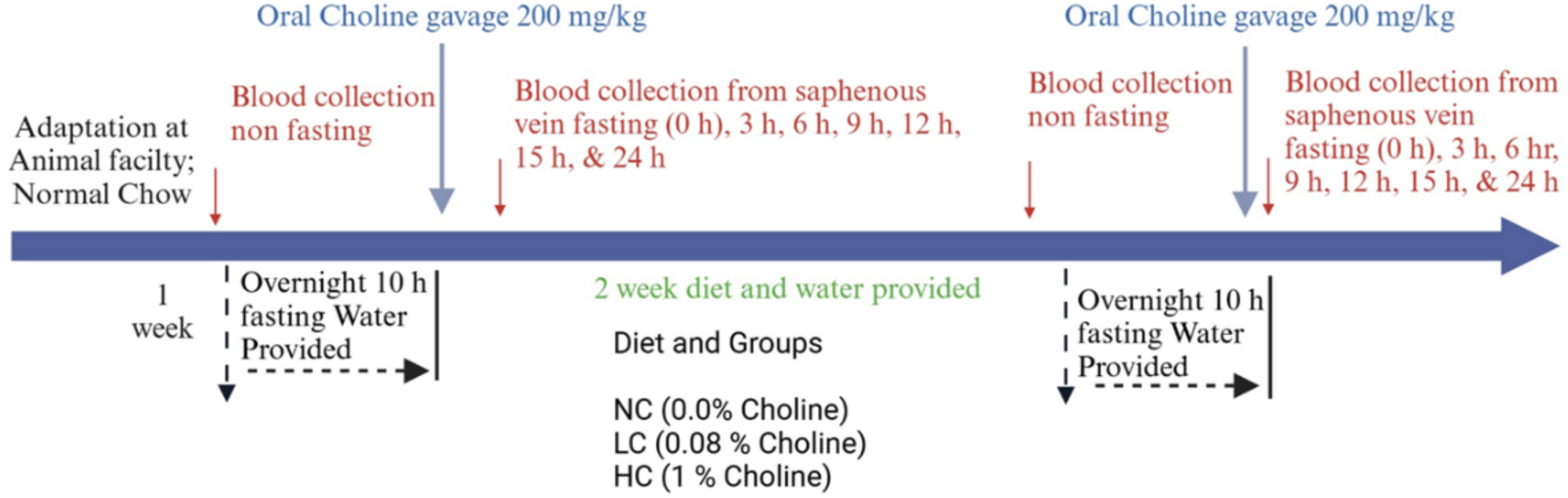
A summary of the experimental timeline and treatments. NC: no choline, LC: low choline, and HC: high choline.

### 2.3 Initial acute choline challenge

Prior to the start of the study, all rats were assimilated for 1 week on the standard chow diet. Non-fasting blood was collected from the saphenous vein in the fed state in the afternoon (5-7 hours prior to fasting) and then rats were fasted overnight for 9-10 h (water provided). The next morning, rats were weighed, and fasting blood was collected from the saphenous vein. Then each rat was orally gavage with choline (200 mg/kg choline, provided as 268 mg/kg choline chloride. Gavage time was noted, and blood was collected at 3, 6, 9, 12, 15 and 24 h after gavage (the fasting blood was time 0 h). Post-gavage, the rats were immediately provided with the chow diet. Immediately following gavage, each rat was observed for dyspnea and signs of fluid in the chest cavity (lethargy, hunched posture, unresponsiveness, changes in normal respiration, etc.) within 0-10 min post-gavage. Rats were observed for signs of distress, such as rough coat appearance, etc, at the end of the day and in the morning of the next day. No animal showed any such adverse signs and symptoms. Blood was directly collected into 0.8 mL lithium Heparin Sep tubes (MiniCollect® Capillary Blood Collection System USA) from saphenous vein using 4 mm lancets. Blood was centrifuged at 3000×g for 10 min, plasma was briefly stored on ice, and then stored at −80°C for further processing.

### 2.4 Dietary intervention

After the first choline challenge was completed (24 h), animals were immediately randomized to three dietary interventions (*n*=4/diet): no choline (NC, AIN-93M mature rodent diet, without added choline, 0% choline bitartrate, D13120601), low-choline (LC, AIN-93M mature rodent diet with 0.08 % choline bitartrate, D10012M), or high-choline (HC, AIN-93M mature rodent diet with 1% choline bitartrate, D230113903) (**Table 1**). The NC diet was designed to eliminate TMAO production from exogenous choline (although TMAO from phospholipids etc. would remain). The HC diet was designed to stimulate chronic TMA/TMAO overproduction based on previous published reports [18,38–41] compared to the LC diet, which was chosen to be not completely depleted of choline but was still less than the 0.2% nutritionally adequate amount commonly used in rodent diets [42,43]. The relevance of these dietary levels to humans is complicated due to distinct nutritional requirements. However, assuming rats eat ∼10% of their body weight per day, the experimental diets corresponded to ∼0, 8 and 100 mg choline/kg body weight/day. Using common body surface area dose conversion formulae [44], these daily doses would correlate to ∼0, 1.3 and 16.2 mg/kg body weight/d in adult humans (or ∼0, 90.8, and 1,135 mg/d in a 70 kg human). Adult human requirements are thought to be 425 and 500 mg/d for women and men, respectively [45]. Thus, the experimental diets provided doses higher and lower than human requirements. All diets (NC, LC, HC) were purchased from Research Diets (RDI, New Brunswick, NJ, USA). Food and water were provided *ab libitum* and monitored regularly.

**Table 1.**
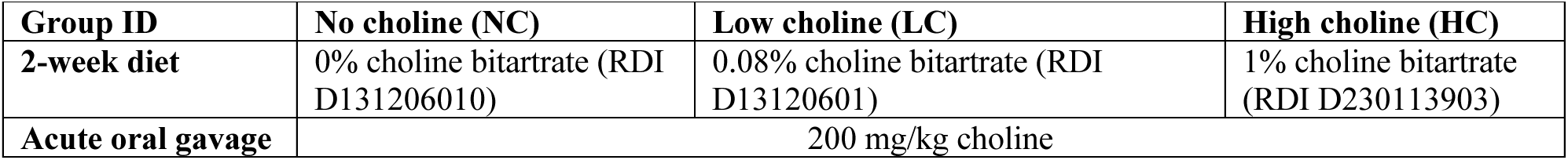
Treatment groups and respective 2-week diet and acute choline challenge parameters.

| Group ID | No choline (NC) | Low choline (LC) | High choline (HC) |
| --- | --- | --- | --- |
| 2-week diet | 0% choline bitartrate (RDI D131206010) | 0.08% choline bitartrate (RDI D13120601) | 1% choline bitartrate (RDI D230113903) |
| Acute oral gavage | 200 mg/kg choline |  |  |

### 2.5 Final acute choline challenge

The same procedures for the first choline challenge (non-fasting blood collection in the afternoon, fasting, gavage, and blood kinetics) were reproduced after the 2-week dietary intervention, except that rats were returned to their respective diets after gavage.

### 2.6 Euthanasia

Animals were euthanized by CO_2_ following current American Veterinary Medical Association guidelines in a closed vessel with CO_2_ from a compressed gas cylinder. CO_2_ flow was controlled to displace 30-70% of the cage volume/min. Animals were exposed to CO_2_ until 2∼3 minutes following apparent clinical death (cessation of respiration). Following euthanasia, the abdominal cavity was opened ventrally from the rectum to the neck, and bilateral pneumothorax was performed to ensure death.

### 2.7 Choline and TMAO extraction and analysis

Choline, and TMAO in plasma samples were quantified by stable-isotope dilution UHPLC-MS/MS as described previously for animal samples [23,46–48]. Plasma samples (25 µl) were plated manually into 96-well plates, followed by automated sample preparation using an OT-2 liquid handler (Opentrons, NY, USA). Acetonitrile (100 µl) and zinc sulfate (10 μL) (5% w/v in water) were added to each well and 20 μL of a mixture of choline-d_9_ and TMAO-d_9_ as internal standards (IS, 2.5 μM each). The 96-well plate was sealed with an Adhesive Plate Seal (Waters, Milford, MA, USA) and sonicated for five minutes at room temperature in a water bath. After sonication, samples were transferred to AcroprepAdv 0.2 µm WWPTFE 96-well filtering plates (Pall Life Sciences, Port Washington, NY) and spin filtered (4 ^0^C, 3400 x g for 10 min) onto Waters Acquity UPLC 700 μL Round 96-well Sample Plates and stored at −80°C until analysis. Choline and TMAO were analyzed by UHPLC-ESI-MS/MS as described previously [30]. Separation was performed on a Waters Acquity UPLC system equipped with an ACQUITY BEH HILIC column (1.7 µm, 2.1 × 100 mm) and pre-column (1.7 µm, 2.1 × 5 mm). The mobile phases consisted of 15 mM ammonium formate (pH 3.5) in water (A) and acetonitrile (B), with an isocratic gradient (80% B) at 0.65 mL/min. Column and autosampler temperatures were 30 °C and 10 °C, respectively. Quantification utilized a Waters Acquity triple quadrupole (TQD) mass spectrometer, operating under positive-mode ESI, with source and capillary temperatures at 150°C and 400°C. The capillary voltage was set at +0.60 kV, and nitrogen flows for desolvation and cone gas were 800 and 20 L/h, respectively. Data acquisition was performed using multi-reaction monitoring (MRM) in MS/MS mode. MRM fragmentation conditions for analytes and IS compounds are reported in **Table 2**.

**Table 2.** Optimized multi-reaction monitoring conditions for detection of choline and TMAO, and their internal standards.

|  |  |  |  |  |  |
| --- | --- | --- | --- | --- | --- |
| **Compound** | **MW** | **RT (min)** | **MS/MS transition** | **Cone Voltage (V)** | **Collision Energy (eV)** |
| Choline | 104.2 | 1.05 | 104.2→60.0 | 38 | 16 |
| Choline-d<sub>9</sub> | 113.2 | 1.05 | 113.3→69.1 | 40 | 16 |
| TMAO | 75.11 | 1.25 | 76.16→58.91 | 40 | 10 |
| TMAO-d<sub>9</sub> | 84.17 | 1.20 | 85.21→68.10 | 40 | 12 |
| MW, molecular weight; RT, retention time, CV, cone voltage; CE, collision energy |  |  |  |  |  |

For sample quantification, LCMS-grade water was spiked with 24 different concentrations of choline and TMAO standards to obtain external calibration curves. Standard samples (25 μL) were then prepared identically to blood samples. Samples were quantified by calculating the analyte/IS peak abundance ratios and then interpolating the concentrations from the standard curves. Data acquisition was carried out using Masslynx software (V4.1 version, Waters).

### 2.8 Data analysis and statistics

Kinetics (concentration vs. time) were plotted and area-under-the-curve (AUC) values were calculated for choline and TMAO from acute challenges (pre- and post-dietary challenge). Values for each analyte were compared by 2-way repeated measures (RM) ANOVA (to test effects of time, treatment, and their interactions) and appropriate *post hoc* tests between treatments. Fasting and non-fasting blood levels and kinetic AUCs were compared pre- and post-dietary choline intervention (within treatments) to assess adaptation over time, and between treatments (post-dietary choline intervention). To assess the effects of non-fasting and fasting between different dietary conditions, i.e., before diet and after comparison on choline utilization and TMAO production, a two-way ANOVA followed by an appropriate test for multiple comparison was used.

## 3. Results

### 3.1 Choline blood concentrations in fasting and non-fasting conditions

Blood concentrations of TMAO is assessed in humans and rodents as biomarkers of CVD risk. However, there is no consensus on whether fasting- or fed-state blood levels or kinetics in response to an acute challenge should be measured, or which is most useful. Therefore, we first assessed fed and fasted blood levels before and after a 2-week dietary intervention with various levels of choline in the diet.

Plasma choline levels in a non-fasting state before the dietary choline had no significant difference between groups as expected due to all groups being assimilated on the same standard lab chow rodent diet (**Fig 2A**, blue color bars). After the dietary choline intervention, non-fasting choline levels were higher in the HC group, followed by LC and NC (**Fig 2A**, red color bars). When comparing non-fasting choline within each diet group before and after the intervention, no difference was observed for either NC or LC groups before and after the diet, while a significant difference was observed in the HC group before vs. after the diet (**Fig 2A**). It is interesting to note that the fasting choline blood levels of the NC and LC groups were not significantly different after going from the chow diet (0.253% choline) both the NC diet (0% choline) or LC diet (0.08% choline). Taken together, these data suggest that large differences in dietary choline levels are reflected in non-fasting blood samples for very high dietary choline. However, at lower dietary levels, some differences are not reflected in non-fasting levels, suggesting somewhat tight control (as the dietary choline in the LC and HC groups differed by 12.5-fold, but non-fasting blood choline was only 2-fold different).

**Figure 2.**
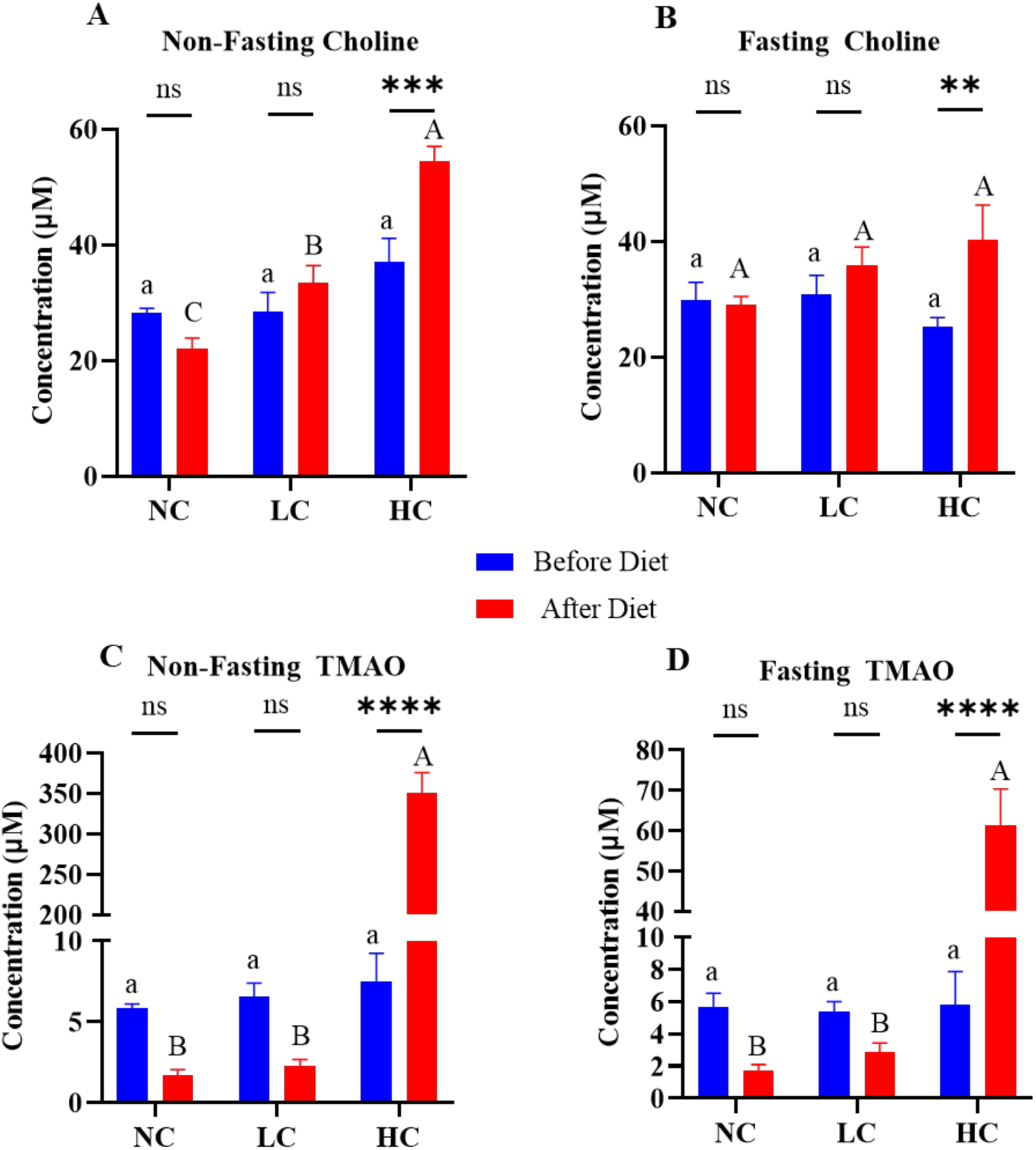
Non-fasting and fasting plasma concentrations of choline and TMAO. **A)** Non-fasting plasma choline before and after dietary choline. B). Fasting choline before and after dietary choline. **C)** Non-fasting TMAO before and after dietary choline. **D)** Fasting TMAO before and after dietary choline intervention. All graphs are (n=4/group). Values are mean ± SEM. Significant differences within diet groups before and after dietary choline are shown by * (ns= no significant difference (p>0.05), * p≤ 0.05, ** p ≤ 0.01, *** p ≤ 0.001, **** p≤ 0.0001); while the difference in various dietary groups before diet and after diet is shown by letters above the graph. Lowercase letters indicate statistical differences between groups before the dietary intervention, uppercase letters represent statistical differences between the groups after dietary intervention; bars sharing letters are not significantly different.

Fasting plasma choline levels are shown in **Fig 2B**. No differences were observed between the 3 diets before or after the intervention. There were also no significant differences within the NC and LC diets between time points. For the HC diet, the plasma choline levels significantly increased after the diet (however, the pre-diet fasting levels were lowest for HC, albeit not significantly lower than NC or HC). This suggests again that circulating choline levels are under tight control, and that fasting levels may not be useful reflections of dietary choline.

### 3.2 TMAO blood concentrations in fasting and non-fasting conditions

Non-fasting levels of TMAO before and after the dietary intervention are shown in **Fig. 2C**. Before the dietary choline intervention, there were no differences between groups (blue bars), again as expected due to all groups being assimilated on the same standard lab chow rodent diet. However, after the diet, TMAO levels for the HC diet were significantly greater than the LC or NC diets (red bars). Within each diet, only HC was significantly different between pre- and post-diet time points (increasing by ∼50x from pre-diet levels). Interestingly, LC and NC levels decreased after the diet, albeit non-significantly compared to baseline. This is because the lab chow diets fed during assimilation was 0.253% choline, which is higher than both NC (0%) and LC (0.08%) diets. These data suggest that high (but not normal or moderate) levels of dietary choline massively increase non-fasting blood levels. Furthermore, increases in TMAO were greatly disproportionate to dietary choline increases: in the HC diet, non-fasting TMAO increased∼50x from pre-diet levels, despite the dietary choline levels only rising ∼4x, from 0.253% in the chow diet to 1% in the HC diet).

Fasting plasma TMAO levels are shown in **Fig. 2D**. Note the distinct scales between **Figs. 2C** and **2D**. Fasting levels showed similar trends as non-fasting levels. Before the diet choline intervention, there were no differences in TMAO between the 3 diets as anticipated (blue bars). After the diet intervention, the blood TMAO for the HC diet was significantly greater than the LC or NC diets (red bars). Within each diet, only HC was significantly different between pre- and post-diet time points (increasing by ∼10x from pre-diet levels, despite only an ∼ 4X increase in dietary choline, from 0.025% to 1%). Comparing non-fasting (**Fig. 2C**) with fasting (**Fig. 2D**) levels, the TMAO values were roughly the same, except for the HC levels post-diet, for which on-fasting levels were ∼350 μM and fasting levels were ∼60 μM. This suggests that high dietary choline significantly increases both non-fasting and fasting TMAO, but that non-fasting levels drop significantly when fasted. This suggests that non-fasting blood collection may be advantageous for observing the most significant differences between treatment groups, with the caveat that some increased variability might be observed due to differences in time since the last meal (although that was not observed in this case, when rodents eat more frequently).

### 3.3 Acute challenge choline kinetics

Beyond the information provided by fed or fasting levels, an acute choline challenge can be useful to assess choline and TMAO responses in blood to a single, defined choline dose (where metabolic capacity is the only variable, as substrate availability is held constant). The controlled dose and timing of a gavage, and the subsequent plasma response, may provide additional information beyond fed state levels (where the time since the last ingestion of TMAO precursors and their amounts is often unknown) or fasting state levels (which depend heavily on elimination efficiency). Thus, a choline challenge can provide information that may more accurately reflect primarily TMAO production capacity.

For the pre-dietary choline challenge (**Fig. 3A**), choline levels in blood demonstrated that the acute challenge worked as hypothesized to moderately elevate circulating choline. All groups had roughly the same fasting choline (0 h), and increased at time point 3 h, indicating oral acute challenge choline appearing in circulation. All the groups peaked at the highest concentration at time point 3 h, after which choline concentrations declined slightly and steadily, with decline back to baseline levels by 24 h.

**Figure 3.**
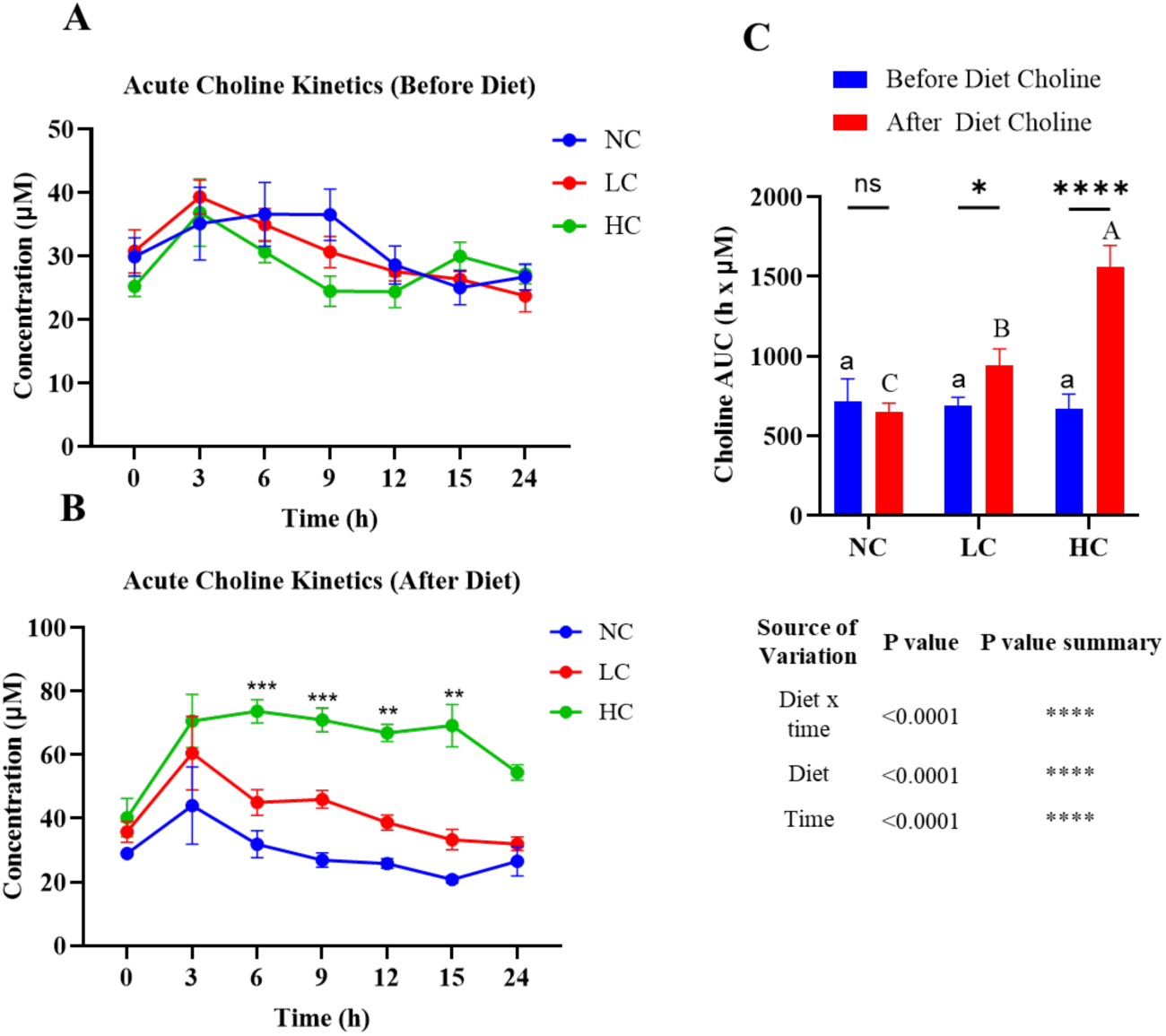
Choline plasma kinetics following acute choline challenge. **A**) Choline kinetics from an acute choline challenge before dietary choline intervention. **B)** Choline kinetics from an acute choline challenge after dietary choline intervention. Data represent the mean ± SEM from n=4 rats/group. The asterisk above at each time point represents significant difference compared to NC (* p≤0.05, ** p ≤0.01, *** p≤0.001). **C)**. Areas under the curve (AUCs) for choline, results presented is a mean of time (h) × Concentration (µM) +SEM (n=4). Lowercase letters indicate statistical differences between groups before the dietary choline intervention. Uppercase letters represent statistical differences between the groups after dietary intervention. Bars sharing the same letters are not significantly different.

In the acute choline challenge after the 2-week dietary intervention (**Fig. 3B**), baseline (0 h) levels were similar between the 3 groups, but the separation was very clear at time point 3 h after receiving the oral gavage of choline. Concentration between the groups and afterwards continues to separate based on various dietary choline availability of either NC, LC or HC. This indicates that even after fasting, the HC group was efficient in maintaining the choline concentration to a considerable level. Also, it’s notable that the choline in NC and LC was very low, and there were slight differences even after the oral gavage of choline both before diet and after diet (Fig **3A**, **3B**). A high choline diet (HC; 1%) effectively alters choline uptake so that serum choline levels rise acutely when given an oral gavage, whereas an NC or LC diet is less effective at increasing choline levels in response to acute administration (**Fig. 3B**).

The areas under the curve (AUCs) for plasma choline kinetics before and after the dietary choline were calculated (**Fig. 3C**). As expected, no differences were found in choline AUCs between the 3 groups before the dietary choline intervention. There was a clear difference observed between all 3 diet groups following the 2-week dietary choline. Within diets, no difference was detected for choline AUCs between the pre- and post-diet intervention challenges for the NC group. Both LC and HC groups showed statistically different AUCs between the pre- and post-diet intervention choline challenges. These data suggest that high dietary choline significantly increases choline plasma concentrations.

### 3.4 TMAO kinetics after acute choline challenge

Plasma TMAO levels were monitored following the acute choline challenges to track TMAO formation and presence in the blood. In the pre-diet choline challenge, similar to the choline kinetics, the TMAO levels dramatically dropped to a base level in all the groups when the rats were fasted (**Fig. 2D**, **Fig. 4A**). After the choline gavage, TMAO levels started to increase slightly at the same rate after the 3 h time point in all the groups (**Fig. 4A**). Maximum levels (∼200 μM) were achieved around 12 h and gradually declined back to near baseline levels around 24 h. Notably, the peak TMAO levels (12 h, **Fig. 4A**) significantly trailed peak choline levels (3 h, **Fig. 3A**), as TMAO formation requires choline to proceed to the lower gut for microbial conversion to TMA and then conversion to TMAO by FMO3 in the liver before appearing in circulation. Also seen for choline (**Fig. 3A**), there were no kinetic differences between groups in TMAO prior to the 2-week dietary choline intervention. For the post-dietary intervention challenge, there was a clear difference in TMAO kinetics between groups, particularly HC (**Fig. 4B**). Also, it was clear that in the after diet regime, the TMAO concentration was higher in the HC group even after the fasting (**Fig. 4B, 2D**), which indicated that 1% dietary choline could be a potential method for future studies, providing very high TMAO acute production which could then be subject to potential inhibitory treatments. Interestingly, the NC and LC groups did not have a major difference in TMAO kinetics after the dietary choline interevntion. For these 2 groups, TMAO increased gradually up to 12-15 h (∼200 μM for NC at 15 h and ∼150 μM for LC at 12 h). However, the HC group started higher and went up to around 350-400 μM at 6 h and remained high through 24 h. These data suggest that high levels of dietary choline significantly elevate fasting as well as postprandial TMAO levels, whereas lower levels (LC, 0.08%) do not.

**Figure 4.**
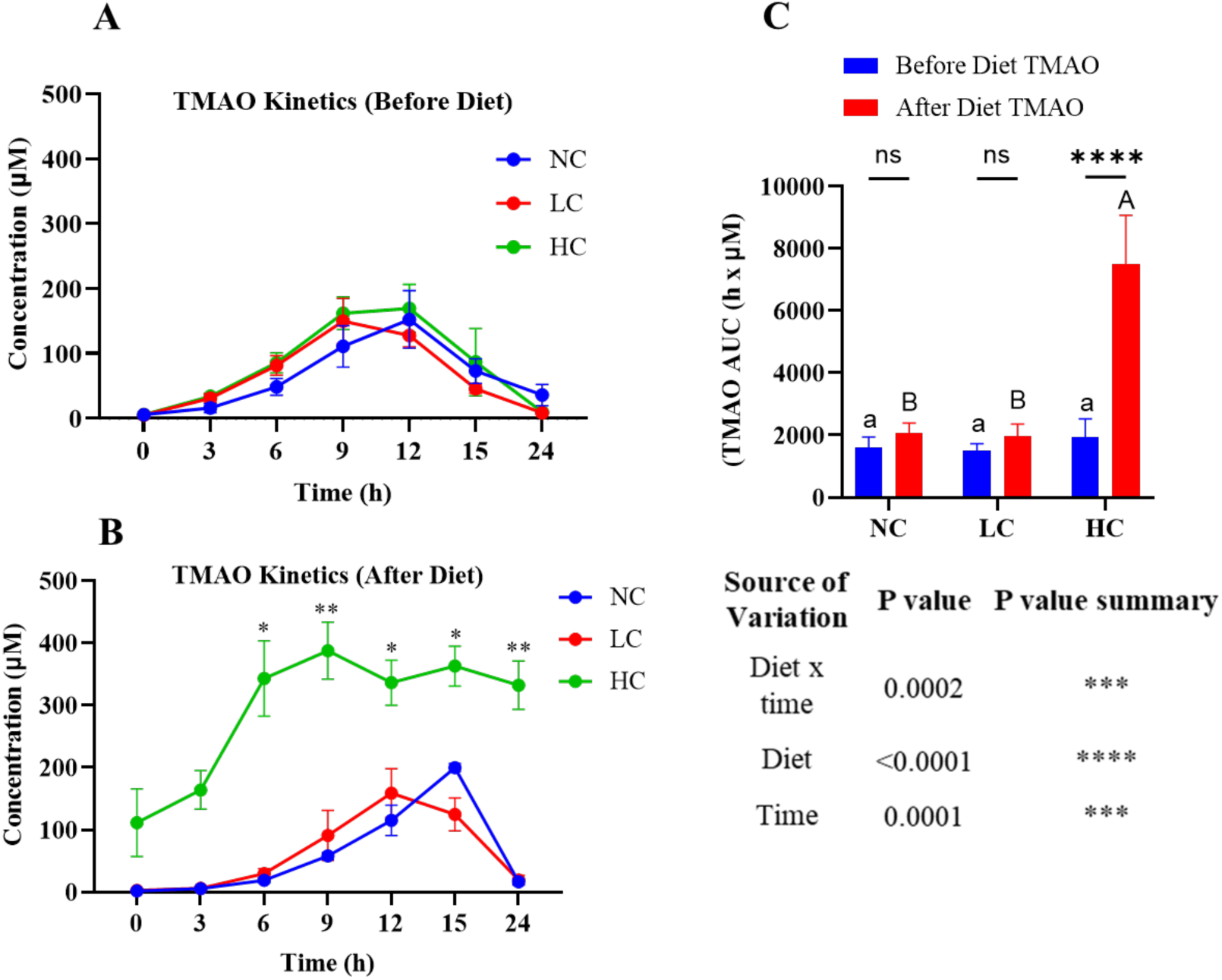
TMAO plasma kinetics following acute choline challenge. **A**) TMAO kinetics from an acute choline challenge before dietary choline intervention. **B)** TMAO from an acute choline challenge after dietary choline intervention. Data represent the mean ± SEM from n=4 rats. The asterisk above at each time point represents significant differences compared to NC (* p≤ 0.05, ** p ≤ 0.01, *** p ≤ 0.001). **C)** Area under the curve for TMAO, results presented as mean of time (h) × concentration (µM) +SEM (n=4). Lowercase letters indicate statistical differences between groups before the dietary intervention. Uppercase letters represent statistical differences between groups after dietary choline. Bars sharing the same letters are not significantly different.

Looking at AUCs of TMAO kinetics following the acute choline challenge (**Fig. 4C**) before the dietary choline intervention, there were no differences between groups (blue bars), as expected. After the dietary choline intervention, NC and LC treatments had the same AUCs (which did not differ from pre-intervention levels). The HC diet significantly increased TMAO after the dietary phase, compared to the HC group pre-intervention and compared to the other two groups post-dietary choline intervention. This suggests that HC dietary treatment can be used in future chronic and acute studies to assess the efficacy of potential TMAO-reducing agents (**Fig. 4C**).

## 4. Discussion

In the past few decades, the prevalence of CVD and associated mortality have increased significantly [49]. The ecology and associated metabolome of the gut microbiome has been found to be associated with CVD progression and as a strong contributing factor. Especially microbial metabolites such as TMAO and tryptophan-derived indoxyl sulfate are known prognostic microbial metabolites that lead to various CVD events [50,51]. To date, there is no approved treatment to reduce levels of TMAO to counteract its vascular consequences, such as atherosclerosis, heart failure, and coronary heart events. Researchers are looking for dietary-based approaches with lower or fewer side effects than conventional therapeutic interventions. While reducing TMAO is the goal of many interventions, there is no consensus on whether interventions targeting choline conversion to TMAO can be more rigorously evaluated by acute oral challenges or fasting/fed levels during chronic dietary studies. Further, to the best of our knowledge, there are no data assessing the comparative utility of fasting and non-fasting conditions in dietary and oral gavage in any in *vivo* study. Fasting and non-fasting TMAO levels are commonly used both in human and animal studies. It is worth mentioning that non-fasting represents a state where choline and TMAO are being consumed and produced, respectively, and eliminated. Fasting, on the other hand, represents a state that persists after choline intake, TMAO production, and TMAO elimination. These are distinct biomarkers, and it is unclear which is more important as a risk factor for CVD. In the present study, we evaluated (i) the plasma TMAO responses to acute oral choline gavage compared to a chronic choline diet, (ii) compared plasma choline and TMAO levels in fasting and non-fasting conditions, and (iii) investigated the impact of diets with or without choline on circulating choline and TMAO. We found a clear difference between treatments of fasting, oral gavage, and dietary choline in the preclinical model. This outcome could be considered for future therapeutic studies using either oral gavage or dietary choline. It also provides a basis for using choline as a precursor and helps establish a rationale for testing therapeutic interventions targeting TMAO *in vivo*.

In our study, all groups had similar choline levels in non-fasting conditions before the dietary choline intervention. However, after the dietary choline intervention, a noticeable difference in choline levels was observed in the non-fasting condition. As shown, there is a range based on the type of diet, with the NC group having the lowest choline levels, followed by the LC group, and the HC group showing the highest choline levels (**Fig. 2A**). Non-fasting choline values, as expected, thus reflect differences in dietary choline levels. Previous studies also demonstrated that fasting could be useful in excluding the inter individual variability and to accurately get a basal level for any intervention in place [52]. In our study, we also measured choline in the fasting condition (**Fig. 2B**). Our results show no difference in fasted choline among dietary groups pre- or post-dietary choline intervention, although fasted choline did increase significantly for the HC diet after the 2-week diet (Fig. 2B). A previous study demonstrated that various blood biomarkers can vary and require optimized fasting conditions [53]. While we did not focus specifically on evaluating fasting times in this study, we selected the minimal fasting period commonly used in rats to minimize stress on the animals [54–56]. Our results show that fasted choline is somewhat under tight control, and dietary differences are poorly reflected in fasting choline. Thus, non-fasting choline appears to be the better biomarker, particularly where differences in choline are suspected or intentionally administered.

Dietary choline levels affected plasma TMAO fasting and non-fasting levels much more than plasma choline levels. The 2-week dietary choline intervention resulted in a massive increase in non-fasting TMAO for HC, while NC and LC were not different (**Fig. 2C**). This suggests that at some level, TMAO production is kept in check by TMAO elimination, but this process is significantly overwhelmed at higher TMAO production levels (a 4-fold increase in dietary choline resulted in a 50-fold increase in non-fasting TMAO levels for the HC group). A similar trend was observed in fasting TMAO levels after the dietary choline intervention (**Fig. 2D**), but fasting TMAO levels in the HC group were reduced ∼5x (from 450 to 70 μM) from non-fasting levels. These data suggest that both non-fasting and fasting TMAO levels are useful biomarkers of dietary choline and conversion to TMAO. However, fasting levels will show lower separation among treatments. Non-fasting TMAO may be the most sensitive and discriminating treatment differentiator.

While fasting and non-fasting markers are useful, the drawback of these methods is that the choline intake amount and time since the last choline intake are not as finely controlled. Additionally, these markers cannot be utilized to determine if a potential TMAO-lowering treatment acts via acute vs. chronic mechanisms. By controlling the timing and amount of choline administered and comparing post-administration kinetics, a more refined biomarker is obtained. Additionally, acute choline challenges administered simultaneously with a potential TMAO-lowering can identify acute inhibitors (which require the inhibitor and substrate to be present at the same time) without prior administration of the treatment of interest. Compounds that act via chronic mechanisms (including shifts in microbiome taxonomic distribution, altered levels of *cutC/D gene copy or CutC/D protein*, and/or changes to FMO3 expression) require an adaptation period and also do not require the TMAO-lowering agent to be present. Thus, with no prior treatment, chronic acting agents would theoretically not inhibit acute TMAO production because they are not CutC/D inhibitors. For this reason, we explored the choline and TMAO kinetics in blood following an acute choline challenge.

In the acute choline challenge, our initial results from before the diet showed that gavage-administered choline entered the bloodstream, reaching a certain level that was maintained for a few hours. The concentration peaked at the 3-hour time point (**Fig. 3A**), aligning with findings from a related previous study conducted in mice, although with different objectives than ours [57,58]. Dietary choline clearly altered the acute kinetics for choline (**Fig. 3B**). Specifically, the HC group sustained a significantly higher choline concentration, as the diet contained added choline. In contrast, choline levels in the LC and NC groups began to decline after the 3-hour point. This demonstrated that a 1% (HC) dietary choline level is effective in maintaining substantial choline levels in circulation.

TMAO kinetics proved highly informative. In the pre-diet challenge, a slight delay was observed, with TMAO beginning to appear and gradually increasing between 3-6 h (**Fig. 4A**). Interestingly, pre-diet TMAO levels in all groups gradually increased. After 9-12 hours, TMAO leveled off and began to decline. This suggests that orally gavage choline offer a narrow but reliably steady time window. Specifically, the 6-12 h period seems optimal for focused analysis, providing valuable insights into the metabolic responses to oral choline intake. However, this limited timeframe presents challenges for conducting therapeutic intervention studies due to its practical constraints. In the after-diet kinetics of TMAO, a clear variation was observed across treatment groups, particularly in the HC group. Unlike the LC and NC groups, which did not appear to exhibit changes in TMAO kinetics after the 2-week dietary choline intervention compared to the pre-diet intervention, the HC group demonstrated a rapid and significant increase in TMAO, with a steady TMAO level after 12 hours. This suggests that dietary choline has a significant impact on TMAO levels, making it particularly useful in studies focused on this metabolite. The difference in TMAO levels in the after-diet condition can likely be attributed to changes in microbiome *cutC/D* copy number, CutC/D protein expression, or gerater activity of *cutC/D*-bearing genera caused by the dietary choline intake [59], and/or increased FMO3 expression.

The type of inhibition studied—whether lowering of fed/fasted levels versus blunting of acute kinetics—depends on the mechanism or proposed action of the inhibitor or TMAO-lowering agent. Interventions that target direct enzyme inhibition of CutC/D or FMO3, for instance, require both the substrate (choline or TMA, respectively) and the prospective inhibitor to be present simultaneously in the gut or liver. These mechanisms are best studied using acute challenges, where the inhibitor is co-administered with choline via gavage. In contrast, treatments that reduce TMAO through changes in microbial composition (e.g., decreasing *cutC/D*-bearing bacteria), microbial function (e.g., reduced CutC/D expression), or hepatic function (e.g., reduced FMO3 expression) are better suited to chronic treatments. These may involve monitoring both fed and fasted states and conducting acute challenges to fully assess the intervention’s effects. The present data suggest that fasting and fed levels are useful for assessing overall TMAO exposure, whereas acute challenges may be useful for parsing out mechanisms of action.

This study is not without limitations. First, we focused on choline and did not include an analysis of carnitine, betaine, or other related precursor molecules of trimethylamine N-oxide (TMAO). The findings and conclusions drawn from this research, therefore, are only applicable to choline. Second, TMA acts as an intermediate metabolite that undergoes conversion into TMAO in the liver before it enters general circulation. Consequently, our study did not include direct measurement of TMA levels due to the inherent challenges involved in such an assessment. The difficulty arises from the rapid and extensive conversion of TMA to TMAO in the liver, making it impractical to accurately capture TMA concentrations. Third, we allowed rats to re-feed on their respective experimental diets immediately following the second challenge instead of reverting to a standard chow diet (which was done for the first acute challenge). This methodological choice may have contributed to differences in the observed kinetics and area under the curve (AUC) measurements. These variations could potentially be attributed to the consumption of the diets with differing nutrient levels post-gavage, rather than to differences in the absorption of TMA or the production of TMAO. Finally, while it would be preferable to use both male and female subjects to ensure a comprehensive analysis, this is complicated by the fact that males are not ideal models for TMAO research due to their low expression of flavin-containing monooxygenase 3 (FMO3), as supported by prior studies[34]. The inclusion of male subjects would require the use of a larger number of animals to compensate for the variability and potentially unreliable data, which may not be a practical or efficient approach [34,35,60–62]. Using twice as many animals for questionable data is not practical. Finally, use of unlabeled choline reduced the specificity of the blood levels of choline and TMAO, as choline from other sources (phospholipids, etc.) could be converted into free choline and TMAO. Use of labeled choline (such as choline-d_9_) in the diet and measurement of the labeled substrate and products (such as choline-d_9_ and TMAO-d_9_) would increase the specificity and sensitivity of these blood biomarkers in animals. Another limitation to this study is that the LC diet (0.08% choline) was actually a moderate choline deprivation diet compared to standard nutritionally complete rodent diets (typically 0.2%). Thus, LC does not represent “normal” levels, but rather lower than sufficient levels. Furthermore, there is a broad gap between LC and HC (1%) diets. Further studies are needed to evaluate normal choline levels (0.2%) as well as intermediately high levels (0.4, 0.6, 0.8%) in order to determine the sensitivity of TMAO response to dietary choline. Also, examination of the impact of dietary choline on tissue levels warrants further study. Finally, extended fasting can be stressful to rodents [63,64]. Our objective for the fast was to get as close to a non-fed baseline state as possible to observe the effects of the acute choline challenge independently from habitual diet. Future dosing studies should investigate the possibility of shorter fasts, as well as using labeled substrates as discussed above to minimize background effects.

It would be ideal to include specific atherosclerosis-related parameters and microbiome diversity analysis in future studies. This approach could provide valuable insights into how the microbiome shifts in response to varying dietary concentrations of specific metabolites. Including both fasting and non-fasting periods in dietary and oral gavage regimens is crucial. This dual approach could help clarify the reasons behind differing TMAO concentrations observed under various conditions. Understanding these variations could reveal how metabolic responses are influenced by different dietary strategies. Such analyses may shed light on the relationship between choline intake, microbiome changes, and TMAO levels. Overall, this could enhance our comprehension of dietary impacts on atherosclerosis risk and metabolic health.

## 5. Conclusions

The outcomes of this study highlighted the importance of differentiating between fasting and non-fasting levels when studying TMAO. The inclusion of a fasting period establishes a baseline that enables a more accurate comparison of subsequent results. However, the non-fasting samples may indicate greater differences between treatments and be more representative of typical blood profiles. Moreover, our findings indicated that administering choline via oral gavage can be used to assess TMAO kinetics. Using dietary choline—particularly at high concentration such as 1% choline—may be the most suitable approach for studies aiming to explore therapeutic strategies involving TMAO. The HC diet appears to provide a consistent and reliable means of raising TMAO levels, which can be beneficial for long-term research into metabolic and cardiovascular outcomes associated with TMAO. However, if the primary research focus involves the study of acute inhibitors of TMAO production or metabolism, the oral gavage method presents a distinct advantage. This approach facilitates the immediate ingestion of choline, followed by rapid absorption and metabolism into TMA and subsequently into TMAO. Such a setup allows researchers to monitor the immediate effects of choline and any interventions on TMAO synthesis, making it an ideal procedure for acute-phase studies where time-sensitive responses are critical.

## Abbreviations

DMB: 3,3-dimethyl-1-butanol
AUC: area-under-the-curve
CVD: cardiovascular disease
CGA: chlorogenic acid
FMO3: flavin-containing monooxygenase 3
HC: high-choline
LC: low-choline
NC: no choline
RM: repeated measures
TMA: trimethylamine
TMAO.: trimethylamine N-oxide

## Author Contributions

Conceptualization: APN, AD; methodology: APN, AD; formal analysis, AD; investigation, AD, AMM, MGS, JGR; resources: APN; data curation, AD; writing—original draft preparation, AD; writing—review and editing, APN, AMM, JGR, MGS, ABPV; visualization, AD; supervision, APN; project administration, APN; funding acquisition, APN, ABPV. All authors have read and agreed to the published version of the manuscript.

## Institutional Review Board Statement

All animal experiments and procedures were approved by The North Carolina Research Campus Institutional Animal Care and Use Committee (IACUC) under approval #20-017.

## Acknowledgments

The authors wish to acknowledge the contributions of Dr. Lisard Iglesias-Carres, who contributed to the original ideation of the study.

